# Accurate in silico predictions of modified RNA interactions to a prototypical RNA-binding protein with λ-dynamics

**DOI:** 10.1101/2024.12.10.627848

**Authors:** Murphy Angelo, Yash Bhargava, Elzbieta Kierzek, Ryszard Kierzek, Ryan L. Hayes, Wen Zhang, Jonah Z. Vilseck, Scott Takeo Aoki

## Abstract

RNA-binding proteins shape biology through their widespread functions in RNA biochemistry. Their function requires the recognition of specific RNA motifs for targeted binding. These RNA binding elements can be composed of both unmodified and chemically modified RNAs, of which over 170 chemical modifications have been identified in biology. Unmodified RNA sequence preferences for RNA-binding proteins have been widely studied, with numerous methods available to identify their preferred sequence motifs. However, only a few techniques can detect preferred RNA modifications, and no current method can comprehensively screen the vast array of hundreds of natural RNA modifications. Prior work demonstrated that λ-dynamics is an accurate in silico method to predict RNA base binding preferences of an RNA-binding antibody. This work extends that effort by using λ-dynamics to predict unmodified and modified RNA binding preferences of human Pumilio, a prototypical RNA binding protein. A library of RNA modifications was screened at eight nucleotide positions along the RNA to identify modifications predicted to affect Pumilio binding. Computed binding affinities were compared with experimental data to reveal high predictive accuracy. In silico force field accuracies were also evaluated between CHARMM and Amber RNA force fields to determine the best parameter set to use in binding calculations. This work demonstrates that λ-dynamics can predict RNA interactions to a bona fide RNA-binding protein without the requirements of chemical reagents or new methods to experimentally test binding at the bench. Advancing in silico methods like λ-dynamics will unlock new frontiers in understanding how RNA modifications shape RNA biochemistry.

## Introduction

Modified RNAs have far-reaching impacts on disease and cellular functions. Over 170 RNA modifications have been identified in biology (1). Many modifications are proposed to be necessary for proper folding of RNA, but some modifications function directly in gene regulation. A classic example is N^6^-methyladenosine (m^6^A). A single methyl group added to the adenosine N^6^ nitrogen leads to the recruitment of the YTH family of RNA-binding proteins, promoting RNA destabilization and turnover (2,3). As a result, the m^6^A modification is proposed to be the single largest determinant of mRNA stability (4) and has disease implications in many types of human cancer and viral pathogenesis (5,6). Thus, RNA modifications can play key roles in biology and pathology through changing how RNA binding proteins interact with RNAs.

RNA-binding proteins interact with mRNA targets to affect mRNA stability and expression. They can bind to any region of an mRNA (7), but RNA binding is critical to exert their functions in transcript regulation (8–10). For example, Pumilio is a prototypical RNA-binding protein and a member of the Pumilio and FBF (PUF) protein family required for embryonic and germ cell development, neuronal differentiation, some human cancers, and other biological functions (11,12). Binding of mRNA by Pumilio is required to recruit other proteins for transcript repression via turnover (11,12). Pumilio binds a conserved UGUAHAUA 8-mer RNA sequence motif typically found in mRNA 3’ UTRs, where H represents A, C, or U (13–18). RNA targets are recognized by a conserved RNA-binding domain consisting of helical PUF repeats (15,19–21). Each PUF repeat discerns a specific RNA base through a three amino acid code (12,22–25). Hence, RNA sequence variations would be expected to affect Pumilio binding, subsequently affecting transcript stability. More recently, two RNA modifications were also noted to change Pumilio-RNA binding affinity in vitro (26), demonstrating that non-canonical modifications can also affect RNA-protein interactions. The impact of the many other known RNA modifications on the binding profiles of RNA binding proteins is still an unresolved question.

Many in vitro and in vivo methods have been developed to identify the preferred binding motifs of RNA-binding proteins. In vitro methods like RNA systematic evolution of ligands by exponential enrichment (SELEX) use recombinant RNA-binding proteins and in vitro selection to identify preferred RNA substrates of a target protein (27). In vivo methods like cross-linking and immunoprecipitation and sequencing (CLIP-seq) enrich for RNA-binding proteins from cell culture or in vivo sources and sequence the associated RNAs to determine the binding protein’s RNA target and footprint (28). Many other strategies and variations of these methods have been reported and are successful in identifying preferred, unmodified RNA sequence motifs. However, none to date can fully identify the effects of the numerous RNA modifications on protein binding. SELEX can screen for preferred interactions with specific RNA modifications (26), but this strategy cannot currently differentiate between modified and unmodified RNA. Mass spectrometry can identify a diversity of RNA modifications (29), but it still relies on RNA sequencing to determine their precise locations within the sequence. Of most concern, a majority of RNA modifications cannot currently be synthesized in vitro. There are a lack of reagents for solid phase chemical RNA synthesis, an inability to incorporate RNA modifications via in vitro transcription, and a lack of methods to add the RNA modifications post synthesis (30). Due to these limitations, current experimental methodologies can study the RNA-protein binding profiles for only a small subset of the 170+ known modifications (30).

Molecular dynamics and free energy calculations are accurate computational methods to model structural complexes and predict the binding affinities of nucleic acid to protein (31–33). Molecular dynamics-based simulations, though time consuming, can illustrate how chemical perturbations affect molecular interactions on an atomistic scale. λ-Dynamics (λD) is an efficient, high-throughput free energy method that can calculate free energy differences corresponding to changes in chemical structure (34,35). As an alchemical free energy method, it uses a sliding λ variable to investigate several structural and chemical modifications simultaneously during a single simulation. After ensuring that all chemical states are sampled equally, λD can calculate changes in binding free energy (ΔΔ*G*_bind_) relative to an original reference molecule. The sign and magnitude of the ΔΔ*G*_bind_ results indicate if a structural modification is favorable or not and by how much. Prior work established λD as an accurate method for predicting relative binding affinities of a library of RNA modifications to modified RNA-targeting antibodies (36). The antibodies in that work were designed to bind singly modified purine RNA bases, and λD accurately predicted off-target antibody binding of other modified bases. Thus, λD demonstrated promise as a practical way to predict the effect of chemical modifications on RNA-protein interactions. That work was limited, however, in that it did not test the accuracy of λD predictions with biologically relevant RNA-binding proteins that bind larger nucleic acid. Establishing such accuracy will be critical to enable comprehensive predictions of RNA-binding protein interactions with λD.

The following study extends prior work testing λD as a viable method to study modified RNA-protein interactions. Pumilio is used as a prototypical RNA binding protein due to its biological and medical relevance, the wide breath of structural and binding studies published, and its large RNA binding footprint. The work further establishes λ-dynamics as a capable method for predicting RNA-protein binding to modified RNAs and identifies computational practices for improved accuracy.

## Results

The goal of this study was to evaluate the accuracy of λD as an in silico strategy for exploring canonical and modified RNA binding to RNA-binding proteins. Pumilio (PUM) was chosen as a prototypical model RNA-binding protein because of its human medical relevance, available high-resolution structures of PUM bound to RNA targets (12,22–25), and extensive studies of its RNA binding specificity (13–18). Like all Pumilio and FBF (PUF) family proteins, human PUM1 (hPUM1) and PUM2 (hPUM2) have a modular architecture of 8 conserved ⍺-helical PUF repeats that facilitates targeting specific RNA sequences (12,22–25) (**Fig. 1A**). Each repeat binds a single base of its RNA target sequence, with a preferred RNA sequence motif of UGUAHAUA, where H is A, C, or U (13–18) (**Fig. 1)**. Prior work determined in vitro protein-RNA binding affinities for a majority of the 4^8^ possible canonical RNA base sequences (18) and for several modified RNAs (26). Thus, hPUM served as an excellent benchmark system for gauging λD’s ability to model protein interactions with both modified and unmodified RNA.

**Fig 1.**
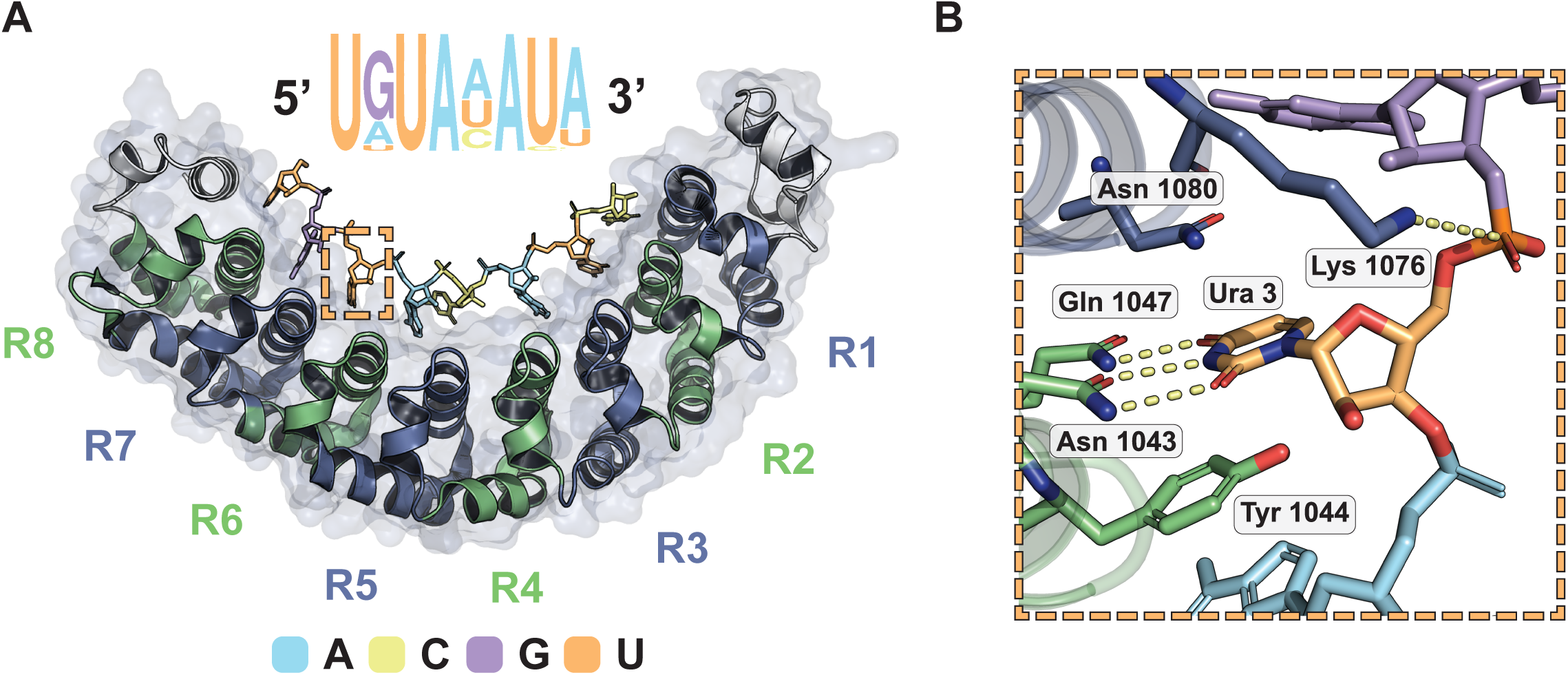
The structure of RNA bound to human Pumilio, a prototypical RNA-binding protein. **(A)** Crystal structure of human Pumilio 1 (PDB ID: 3Q0P) bound to a consensus RNA sequence. The Pumilio RNA binding domain consists of eight helical PUF repeats forming a comma-shaped structure that binds RNA on its interior. Each PUF repeat (R1-R8) interacts with a single nucleobase. Binding is anti-parallel, with the C-terminus of Pumilio interacting with the 5’ end of the RNA sequence motif. Human Pumilio consensus motif presented, with letter size indicating in vivo sequence preference. Dashed box region enlarged. A, Adenine; G, Guanine; C, cytidine; U, uridine. **(B)** Molecular detail of a single PUF repeat (R6) binding to a single nucleobase (uridine). A PUF repeat amino acid triplet binds to preferred bases through hydrogen bonding and stacking. Images by PyMOL.

### λ-Dynamics accurately modeled Pumilio binding to canonical RNA mutants

λD was first used to explore how canonical base mutations were predicted to affect RNA binding to Pumilio. Each of the eight nucleobase sites along the RNA bound to hPUM1 and hPUM2 were mutated to a different canonical base, effectively modeling all possible single nucleotide polymorphisms of the PUM consensus motif. λD calculations were performed with the CHARMM molecular modeling package and the BLaDE GPU accelerated engine in the isothermal-isobaric ensemble (37–39) (see **Methods**). This work expanded upon the procedure used previously to establish λD’s efficacy for modeling antibody binding to ribonucleosides (36). That system used a CHARMM (40) RNA base library for the force fields parameters, the descriptions of the interatom force potentials of a modeled molecule. Other force fields for RNA, like Amber (41), have been used in molecular dynamic simulations but had not been tested with λD. To evaluate the effect of force field accuracy, both CHARMM and Amber-based nucleic acid parameters (39–42) were used in parallel with λD in this work. Computed binding results were then compared to in vitro binding affinities (Δ*G*_bind_) from existing literature (18) (**Fig 2**).

**Fig 2.**
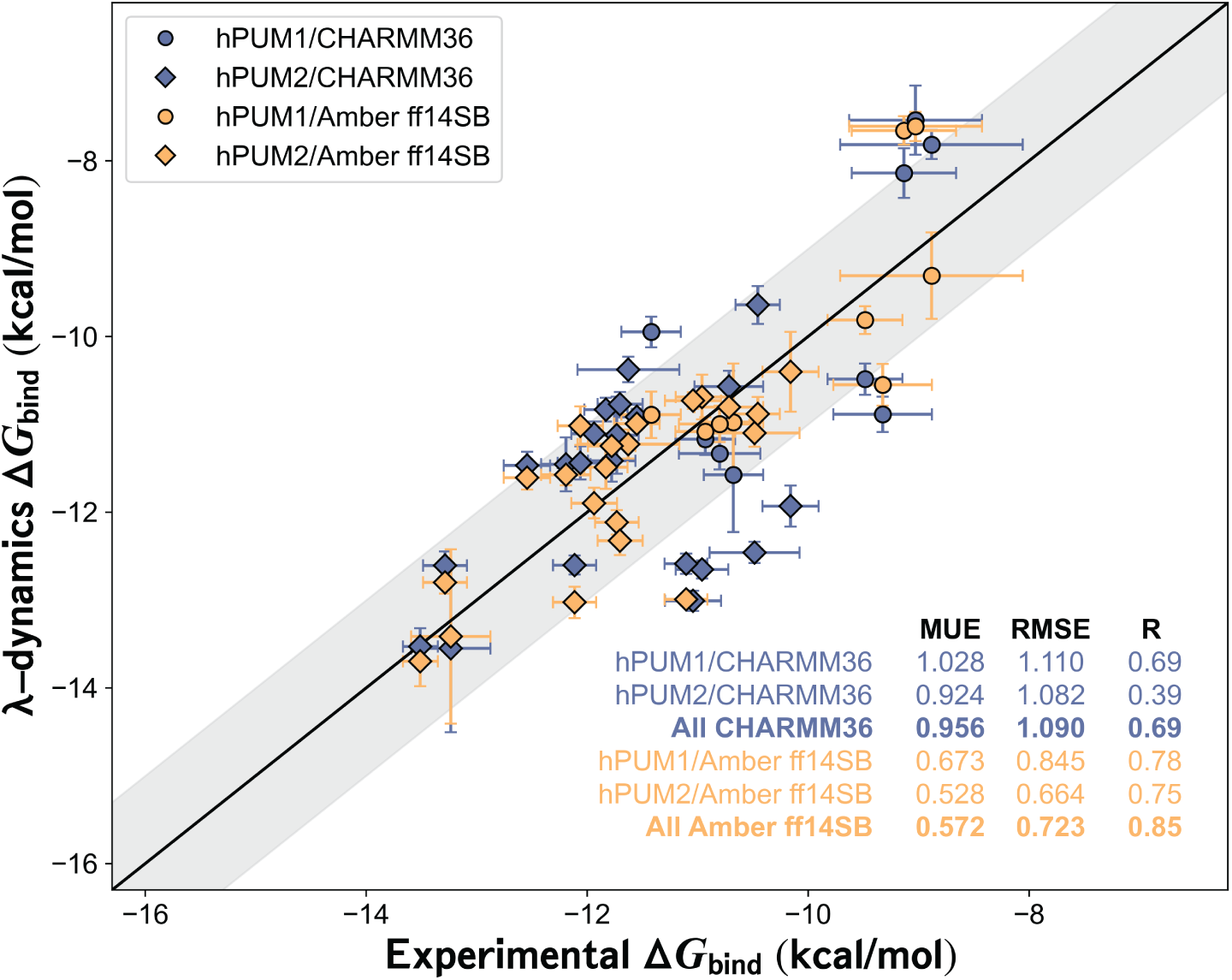
Pumilio-RNA sequence preferences predicted with λ-dynamics. Comparison of in silico λ- dynamics and in vitro measurements of Pumilio binding to a variety of RNA sequences with canonical RNA. Experimental binding affinity results (x-axis) from a previous publication (18). These results were compared against the predicted λ-dynamics Δ*G*_bind_ (y-axis) generated in this study. Both variables reported as kcal/mol. See **Methods** for details regarding the calculations and RNAs selected. Shaded area, root mean square error (RMSE) of 1.0 kcal/mol. Error bars report standard deviation from both data sets. Mean unsigned errors (MUEs) and RMSEs reported for human Pumilio 1 (hPUM1) and human Pumilio 2 (hPUM2) results computed with CHARMM (CHARMM36, blue) and Amber (Amber ff14SB, gold) force fields.

All λD results exhibited strong agreement with in vitro measurements (**Fig. 2**). Mean unsigned errors (MUE) and root mean square errors (RMSE) between the λD predicted free energies differences (Δ*G*_bind_) results and previously published experiments (18) were within or below the accepted gold standard of approximately 1.0 kcal/mol agreement (43–46) (**Fig. 2**, **Table S1**). As a control, each native RNA base was perturbed to an identical but physically distinct copy of itself. These relative binding free energies differences (ΔΔ*G*_bind_) were near zero, as expected of a base replacing itself, indicating that the λD calculations were functioning properly (**Table S1**). Pearson R values indicated strong predictive trends for CHARMM (R = 0.69) and Amber (R = 0.85) force fields (**Fig. 2**). An improvement in accuracy was noted with the Amber ff14sb force field. Relative to CHARMM, MUEs improved by 0.38 kcal/mol, RMSE by 0.37 kcal/mol, and the Pearson R by 0.16 (**Fig. 2**). Terminal RNA bases represented the greatest challenge for λD, with sites 1, 7, and 8 containing predictive outliers exceeding ± 1.0 kcal/mol in each screening (**Table S1**). This may be due to the terminal RNA bases’ ability to rotate out of the PUF pocket and therefore unbind mid-λD simulation. The relatively weaker predictions of the CHARMM versus Amber force field suggested an area of future parameter refinement for modeling RNA interactions with CHARMM. Other factors like slow conformational changes may also have important energetic contributions (see **Discussion**). Regardless, λD was able to accurately estimate hPUM1 and hPUM2 binding trends to canonical RNA. These results also suggest that λD had the potential to predict protein– RNA interactions of expanded chemical complexity.

### λ-Dynamics predicted Pumilio binding to modified RNA

The efficacy of λD was further tested by probing how a library of RNA modifications affected hPUM1 binding. Of the 170 plus RNA modifications currently identified, many do not have commercially available reagents or lack protocols for in vitro syntheses. Following previous work (36), a library of 44 modified nucleobases was selected for λD screening based on the availability of commercial reagents and force field parameters. Using CHARMM force fields (47), λD screened 352 single-site mutants created by applying these 44 modifications at each position along the hPUM1 8-mer RNA consensus motif (see **Methods**, **Fig. 3**). Examples of the results obtained are shown (**Fig. 4**), with full results reported in the Supplement (**Table S2)**. Relative binding ΔΔ*G*_bind_ of the modified RNAs ranged from positive to negative, designating worsened to enhanced binding, respectively, relative to the wildtype RNA sequence. The λD predictions suggested that most modifications negatively impact RNA binding, some have little effect, while others enhance it (**Fig. 4**, **Table S3**). Using a conservative cutoff of ΔΔ*G*_bind_ ≤ –1.0 kcal/mol, approximately a 5-fold or greater enhancement of binding, a total of six modifications markedly enhanced binding to hPUM1 at 3 separate RNA sites (**Fig. 4**, **Table S2**).

**Fig 3.**
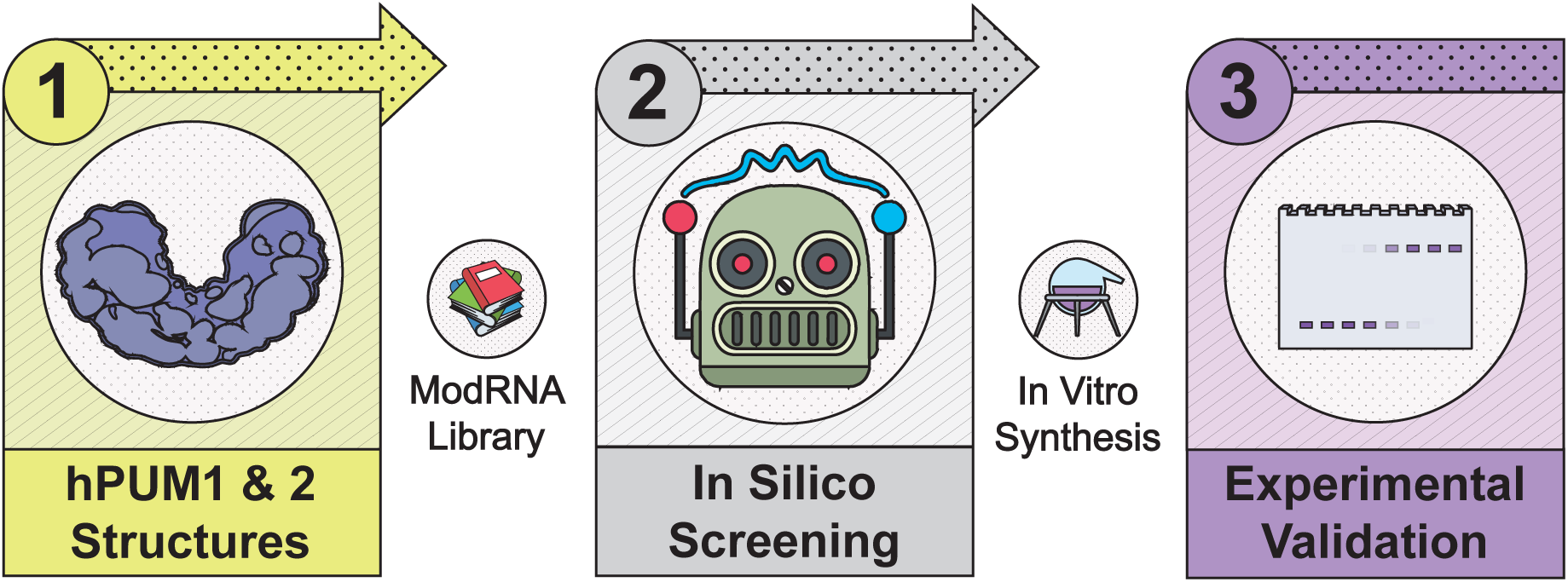
Strategy to test the predictive value of λ-dynamics for modified RNA and RNA binding protein interactions. (1) λ-Dynamics and a modified RNA library were used with high-resolution crystal structures of human Pumilio 1 (hPUM1) and human Pumilio 2 (hPUM2) to predict modified RNA-protein binding. (2) Each RNA position in hPUM1 or hPUM2 were changed to a different RNA and the change in free energy binding was calculated compared to the original RNA-protein structure. See **Methods** for more details. Some of the modified RNA modifications tested in silico could be synthesized in vitro. (3) These modified RNAs were tested for RNA binding to probe λ- dynamics prediction accuracy.

**Fig 4.**
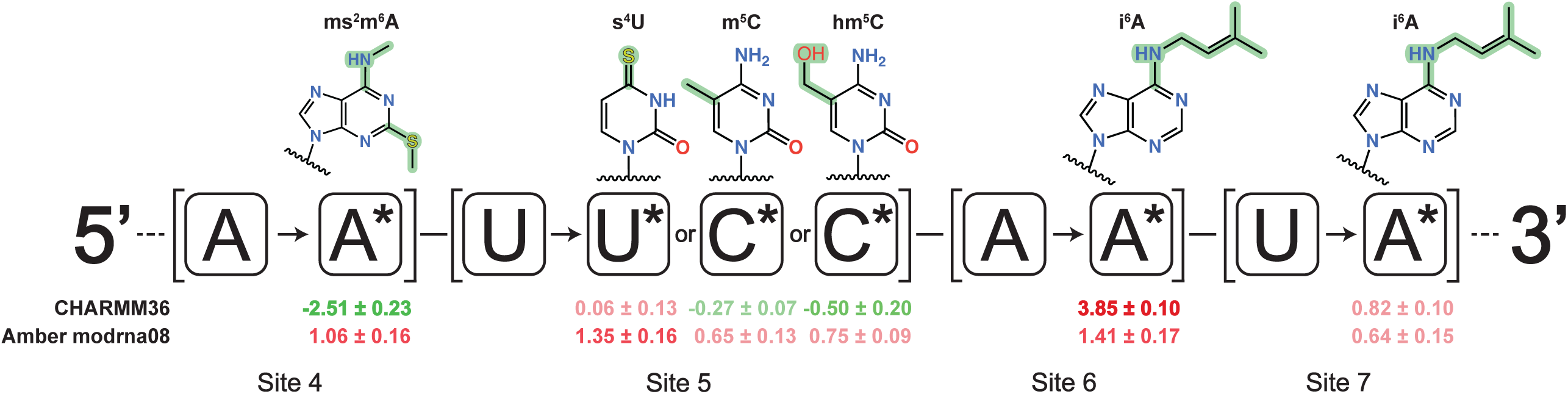
Example λ-dynamics predictions of how RNA modifications may affect binding to human Pumilio 1. The original RNA base is reported, along with the base change (*, RNA name and chemical structure above) and noted relative free energy binding change measured (ΔΔ*G*_bind_, kcal/mol) with CHARMM (CHARMM36) and Amber (modRNA08) force fields. Predicted decrease in binding affinity, positive ΔΔ*G*_bind_, in red. Predicted increase in binding affinity, negative ΔΔ*G*_bind_, in green. Results highlight modifications tested in vitro. Sites refer to RNA positions described in **Fig 1**. See **Table S2** for complete results.

To investigate force field contributions to λD predictions, modified RNAs with worsened, neutral, and enhanced affinities were selected for extended follow-up 50 ns simulations with both CHARMM and Amber force fields (47,48) (see **Methods, Fig. 4**, **Table S3**). Modified RNA selection was also based on the availability of commercial reagents or published in vitro data (26). The extended CHARMM simulations displayed good agreement with the initial, shorter screening. However, the Amber simulations were noticeably different from CHARMM for some of the data points (**Fig. 4**). Three of the Amber predictions differed from their CHARMM counterparts by more than 1.0 kcal/mol (**Fig. 4**). At site 4, 2-methylthio-N6-methyladenosine (ms^2^m^6^A) was predicted to have moderately worsened binding with Amber rather than strongly favorable binding affinity with CHARMM. The Amber vs CHARMM ΔΔ*G*_bind_ differed by more than 3.0 kcal/mol. At site 5, 4-thio- uridine (s^4^U) worsened binding with Amber rather than having no significant change with CHARMM. The Amber vs CHARMM ΔΔ*G*_bind_ differed by approximately 1.3 kcal/mol. These energetic discrepancies between sulfur-containing bases may reflect differences in how each force field treats sulfur-containing moieties, a traditionally difficult element to parameterize accurately with molecular mechanic force fields (49,50). Finally, at site 6, N^6^- isopentenyladenosine (i^6^A) saw an Amber vs CHARMM ΔΔ*G*_bind_ of approximately 2.4 kcal/mol (**Fig. 4**). In summary, λD predicted that several RNA modifications would site-specifically alter hPUM1 RNA binding affinity. Moreover, different RNA force fields yielded different predictions, and thus alternative methods were warranted to determine which computational strategy was more accurate.

### λ-dynamics’ predictions with Amber force fields closely matched Pumilio binding to modified RNA in vitro

To evaluate the accuracy of the λD predictions, computed ΔΔ*G*_bind_ were compared to experimentally determined modified RNA binding by Pumilio, measured here and in prior publications. First, Electrophoretic Mobility Shift Assays (EMSAs) were used to measure the in vitro binding affinity of select modified RNA to hPUM1 and hPUM2. Seven RNA modifications were selected based on commercial availability and the extended screening results (**Fig. 4**). These RNA modifications and paired, unmodified RNA controls were synthesized as 5’- fluorescein-capped RNA oligos through solid-state chemistry (see **Methods**). The modified RNA oligos changed a single site in the UGUACAAU hPUM recognition motif, with unmodified RNA serving as a control (**Fig. 5**). The 5-hydroxymethylcytidine (hm^5^C) building block was only available as DNA and therefore compared to an equivalent DNA-RNA chimera control (**Fig 5**). Fluorescein-labeled RNAs were incubated with increasing concentrations of hPUM1 or hPUM2, then run on a non-denaturing PAGE gel under low voltage to separate unbound versus PUM-bound RNA. The lower and upper bands, corresponding to unbound and bound RNA, respectively, were quantified and dissociation constants (K_d_) were estimated based on the final curve results (see **Methods, Fig. 5B**, statistics in **Fig. S1**).

**Fig 5.**
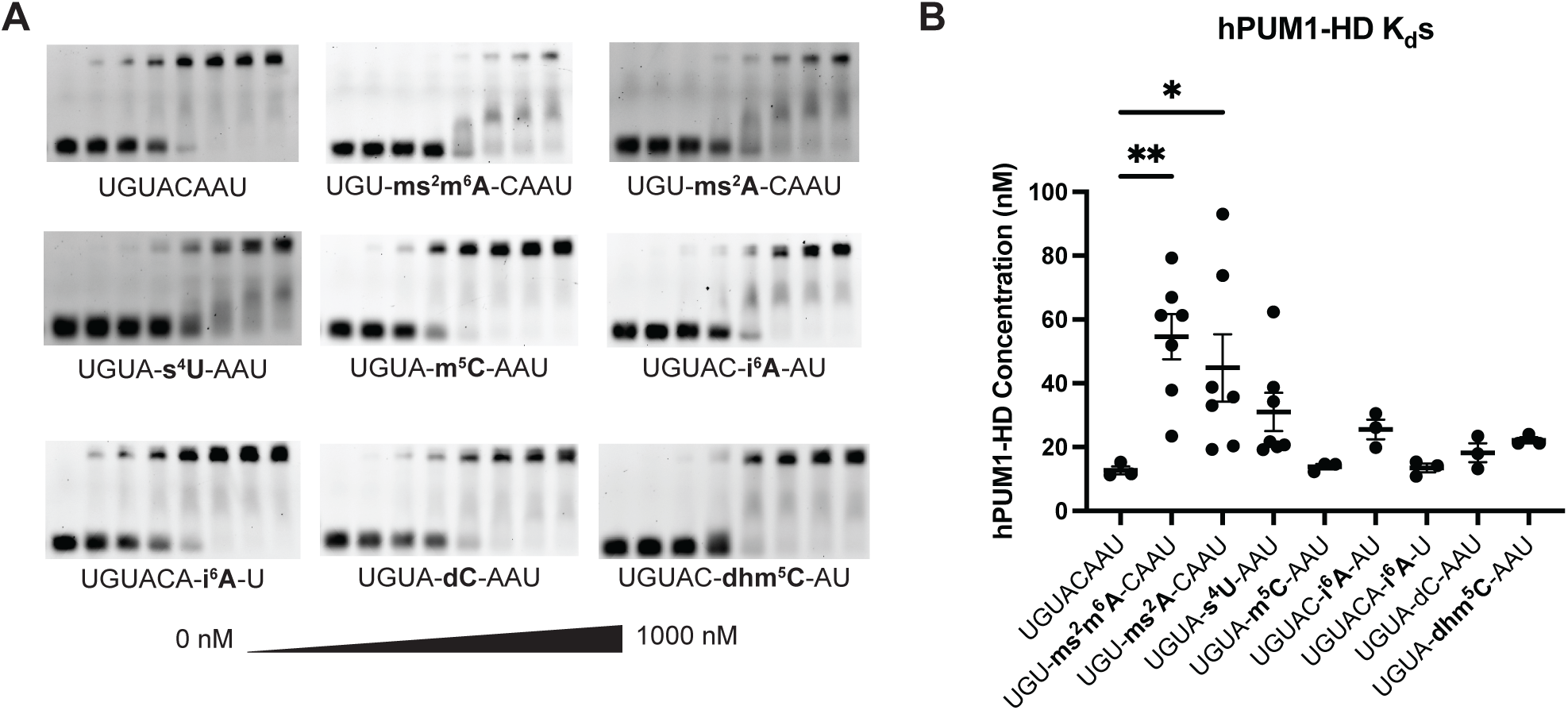
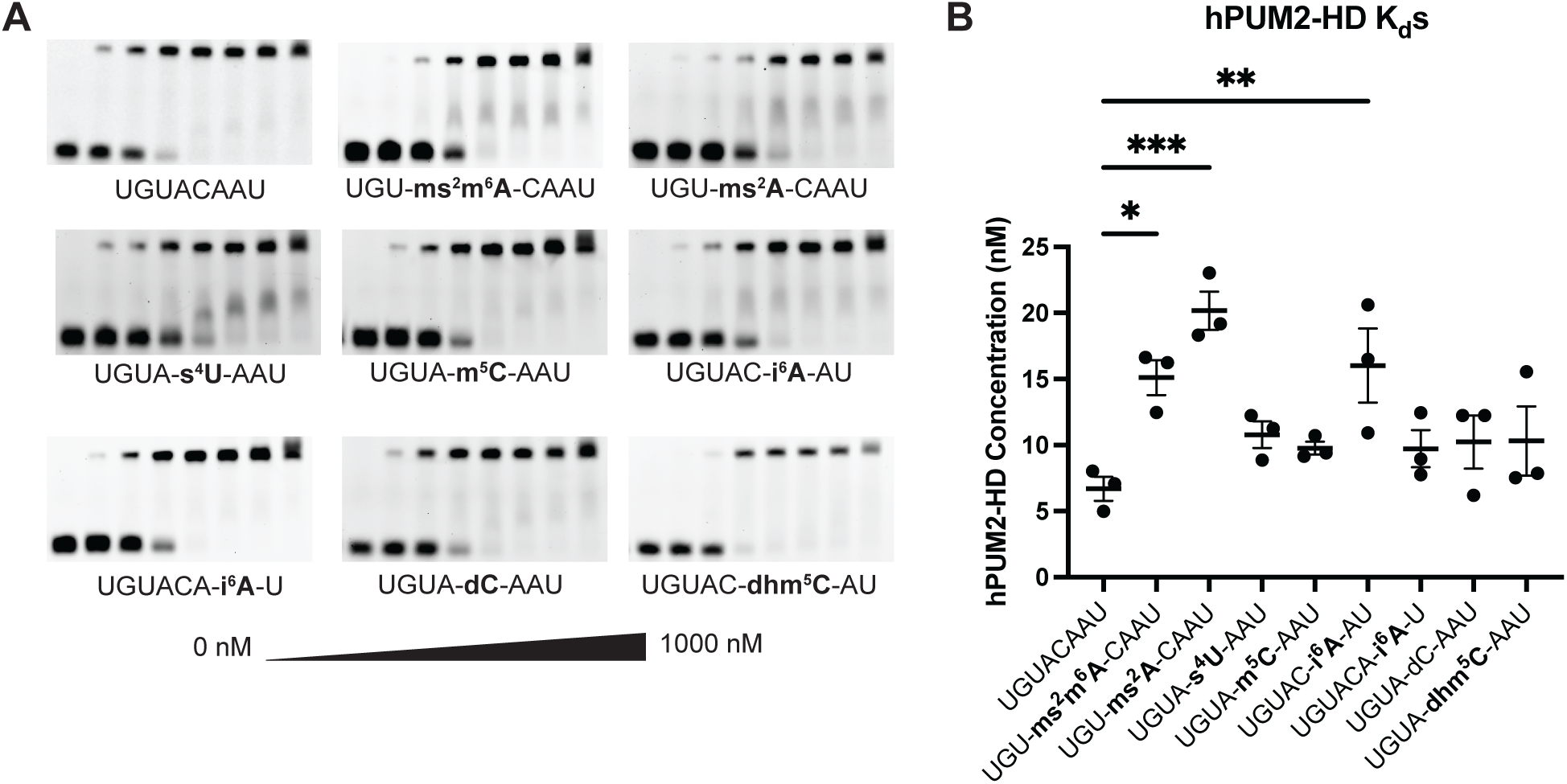
Human Pumilio 1 binding to modified RNAs in vitro was consistent to λ-dynamics predictions. **(A)** Electrophoretic Mobility Shift Assays (EMSAs) were used to estimate the binding affinity in vitro. RNA oligos without or with the designated RNA modifications were incubated with increasing concentrations (0-1000 nM) of recombinant human Pumilio 1 (hPUM1) RNA binding domain, run on a polyacrylamide gel, and imaged for carboxyfluorescein (FAM) fluorescence. A sample without protein served as an unbound RNA control. The lower band corresponds to unbound RNA. The upper band corresponds to RNA bound to the recombinant protein. The binding affinity can be estimated by calculating the protein concentration at the binding inflection point. Shown are representative gels from hPUM1 EMSA binding experiments. **(B)** EMSAs were performed at least three times, and calculated binding dissociation constants (K_d_) reported with their mean and standard deviation. *, p <0.05; **, p <0.01. See **Fig S1** for full statistical analyses and the methods for more experimental details.

The in vitro findings supported the λD predictions and demonstrated that the tested RNA modifications all had unfavorable effects on hPUM1 or hPUM2 binding (**Fig. 5** and **Fig. S2**). With hPUM1, three of the seven modified RNA oligos (hm^5^C and m^5^C at site 5, along with i^6^A at site 7) showed affinities comparable to their respective wildtype. The remaining four (ms^2^m^6^A and ms^2^A at site 4, s^4^U at site 5, and i^6^A at site 6) significantly weakened binding (**Fig. 5B**). Similar results were also observed with hPUM2 (**Fig. S2**). Next, the λD screening results were compared to a prior study measuring Pumilio binding to select RNA modifications. Vaidyanathan, *et al.* measured in vitro binding affinities of hPUM2 to RNA containing pseudouridine (Ψ) or m^6^A RNA modifications (26). That study noted that both modifications weakened RNA binding to Pumilio in vitro as more RNA sites incorporated the modifications. Thus, in vitro binding data supports the binding trends predicted by λD (**Fig. 6**). While no distinctly favorable RNA modifications were identified in this study, several promising candidates remain untested due to lack of chemical reagents, underscoring opportunities for future exploration (see **Discussion**).

**Fig 6.**
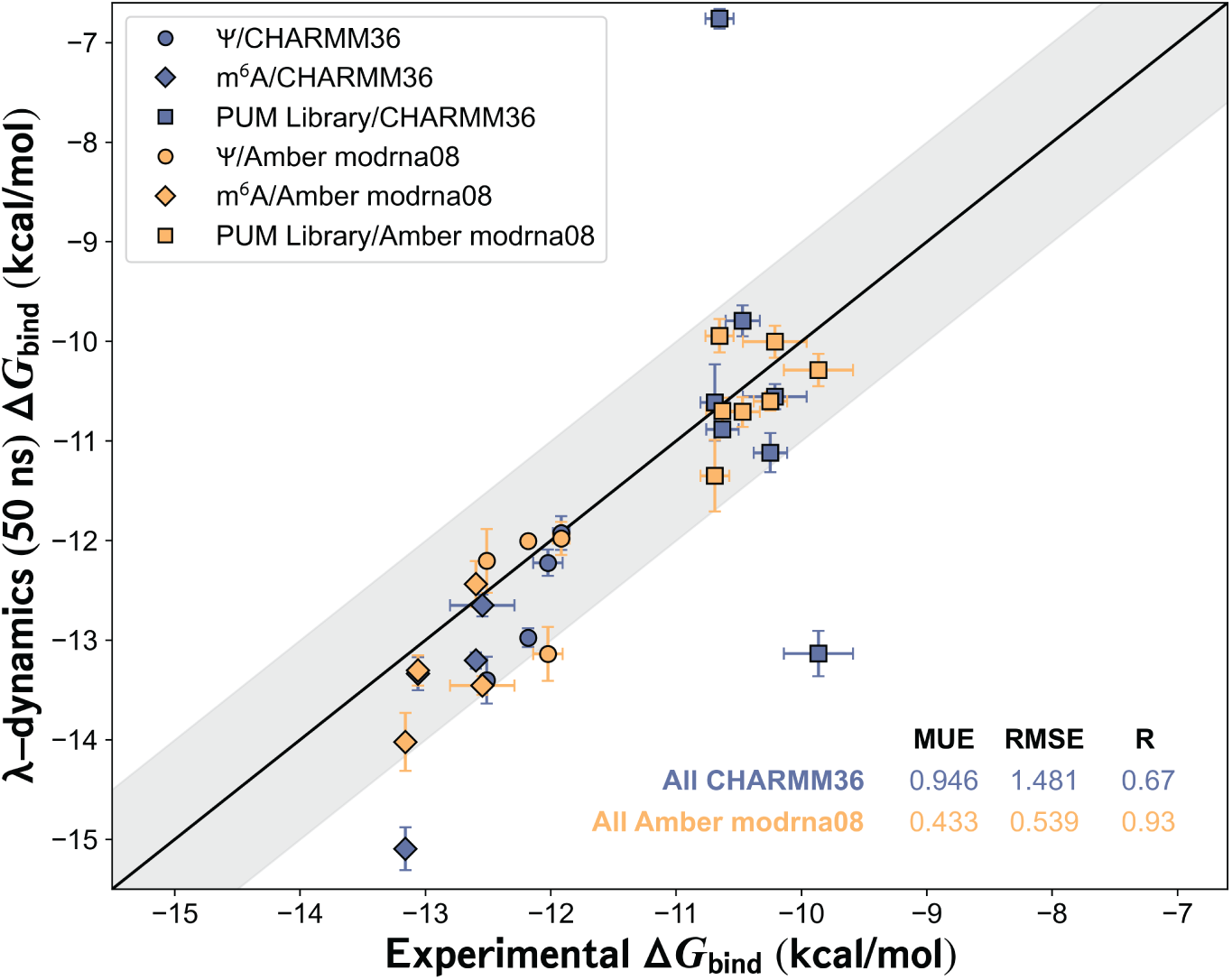
Pumilio-modified RNA interactions can be predicted with λ-dynamics and Amber force fields. Comparison between λ-dynamics and in vitro Δ*G*_bind_ measurements of human Pumilio 2 (hPUM2) binding to pseudouridine (Ψ) or N6-methyladenosine (m^6^A) modified RNA sequences performed previously (26), or of binding data generated in this study (“PUM Library”). The experimental binding data (x-axis) is plotted against the extended 50 ns λ-dynamics simulation results (y-axis) in kcal/mol. See **Methods** for details regarding the calculations performed. Shaded area, root mean square error (RMSE) of 1.0 kcal/mol. Error bars report the standard deviation from both experimental sets. Mean unsigned errors (MUEs) and RMSEs reported for hPUM2 results computed with CHARMM (CHARMM36, blue) and Amber (Amber modRNA08, gold) force fields.

The in silico predictions and binding results were analyzed further to gauge the accuracy of the CHARMM versus Amber force fields (**Fig. 6**). Overall MUEs and RMSEs between λD and experiment were 0.95 and 1.48 kcal/mol for CHARMM and 0.43 and 0.54 kcal/mol for Amber, respectively (**Fig. 6**, **Table S3**). While both force fields demonstrated relatively good predictive accuracy, the CHARMM predictions included three large outliers, resulting in higher average errors (**Fig. 6**). The CHARMM Δ*G*_bind_ for ms^2^m^6^A at site 4 and i^6^A at site 6 deviated from experiment by over 3.0 kcal/mol (**Table S3**). When comparing the binding results of Vaidyanathan, *et al.* (26), one oligo from the m^6^A subset deviated from experiment by almost 2 kcal/mol. The CHARMM predictions achieved a Pearson R of 0.67, indicating moderate predictive capability, but the Amber predictions had a significantly improved Pearson R of 0.93, indicating very strong correlation (**Fig. 6**). The visibly tighter fit of the Amber force field results reflect the improved RMSE compared to CHARMM. Thus, Amber RNA force fields performed more accurately than CHARMM RNA force fields for this prototypical protein-RNA complex. These results collectively demonstrate that the λD in silico method can accurately predict in vitro RNA binding protein interactions with both unmodified and modified RNAs.

## Discussion

Nearly two hundred RNA modifications have been identified in biology, yet there are only a small number of methods to determine how they affect RNA-protein interactions. In vitro methods are limited by the lack of reagents for modified RNA synthesis, and in vivo methods are limited by their inability to detect all modifications within RNA. This work utilized λD free energy calculations to test how a library of RNA modifications could affect single-stranded RNA-protein interactions. Using human Pumilio as a prototypical RNA binding protein, these results demonstrated that λD could accurately predict both unmodified and modified RNA binding without the dependence of in vitro chemical reagents to probe such interactions.

Simulation of biological macromolecules requires an accurate description of their physical properties, such as the force required to stretch or rotate a phosphodiester bond or nonbonded electrostatic and van der Waals interactions. These terms are collectively encoded as force field parameters in molecular mechanics simulations, such as molecular dynamics. Both Amber and CHARMM RNA force fields have been developed to include modified RNA parameters, which have been optimized to accurately model RNA stability and conformational dynamics (39–42,47,48). However, pairing current modified RNA force fields with Amber or CHARMM protein force field counterparts has not been thoroughly investigated. In this work, protein and nucleic acid force fields from Amber and CHARMM were paired to predict changes in binding free energies associated with nucleobase perturbations to canonical and modified RNAs. Notable improvements in predictive accuracy were observed with Amber compared to CHARMM, despite using a slightly older protein force field with Amber (Amber ff14SB) (41,42,48) than with CHARMM36 (39,40,47). Their performance differences further highlight the critical influence of force field selection for free energy calculations. The RNA-PUM results herein suggest that either Amber alone or a CHARMM with Amber consensus approach should be used to maximize predictive accuracy for prospective analysis of RNA-protein binding interactions. Work is ongoing to identify if inaccuracies also exist in Amber RNA force fields for specific chemical moieties not explicitly evaluated in this work. This additional work will be valuable for minimizing false negative and false positive binding predictions with λD.

Understanding how modified RNAs interact with RNA-binding proteins is an active area of investigation. This includes investigating sequence preferences of modified RNA binding proteins and how different RNA modifications affect canonical RNA-binding protein interactions. For example, in vitro SELEX and in vivo CLIP-seq methods have determined that the YTH family of RNA binding proteins have a sequence preference for G(m^6^A)CH (51–53), while a wide variety of methods have determined RNA sequence preferences for human PUM (e.g.,(17,26,54,55)). In vitro RNA binding assays showed how Ψ and m^6^A modifications have a cumulative, negative affect on human PUM-RNA binding (26). In this study, in silico λD correctly predicted many RNA modifications to have negligible or modest effects on binding, consistent with trends observed in vitro. The findings underscore the potentially nuanced and site-dependent effects of RNA modifications on protein binding, a conclusion reached in prior studies (26). Comprehensive screening across all RNA sequence positions and with various RNA modifications is needed to fully understand how RNA modifications influence RNA-protein interactions.

In silico methodologies enable new opportunities to study molecular interactions without the limitations that chemical synthesis or molecular biology requirements place on in vitro or in vivo strategies. This work demonstrates how λD can be used to accurately screen preferred sequence binding motifs for unmodified RNA. More excitingly, the results also support that RNA-binding protein interactions with non-canonical RNAs can be predicted in silico (**Fig. 7**). While most RNA modifications will likely disrupt RNA-protein interactions, a few may enhance binding, leading to breakthroughs in epitranscriptomics and towards the rational design of unnatural, RNA-based therapeutics. With λD and other in silico strategies, probing any RNA-protein interaction is possible.

**Fig 7.**
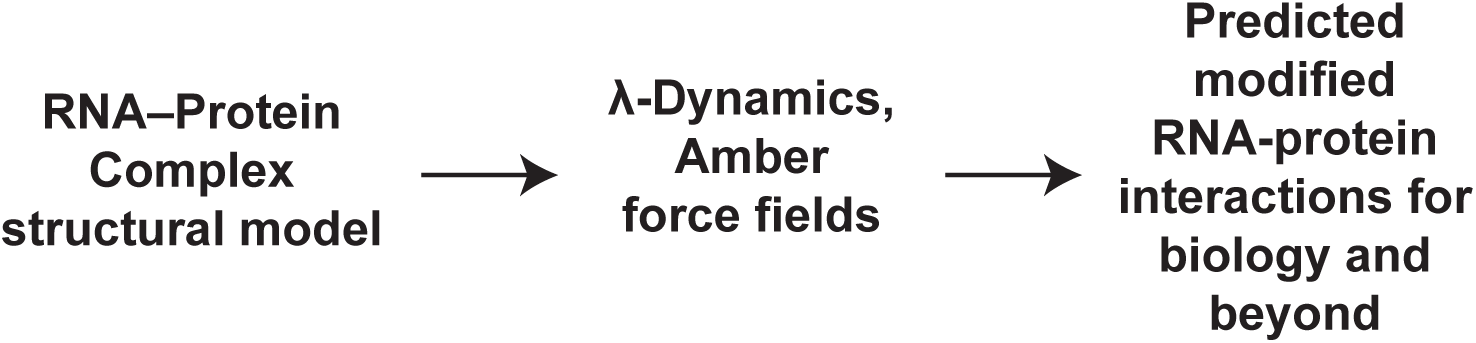
Strategy pairing in silico λ-dynamics with in vitro binding studies to elucidate modified RNA- protein interactions.

## Materials and Methods

### Recombinant protein expression

Human PUM1 (hPUM1) and PUM2 (hPUM2) RNA binding homology domain (HD) coding regions were codon optimized for E. coli, synthesized, and cloned into pET21a or pET28b expression vectors with a His6::V5::PrecissionC protease cleavage site or a His6::SUMO tag, respectively. The plasmids were expressed in LOBSTR cells (Kerafast EC1002), grown in Luria Broth (LB) media until 0.6 – 0.8 OD (600 nm absorbance), then induced with 0.1 mM IPTG and cultured at 16°C and 160 rpm overnight. Bacteria were pelleted and frozen until use. Pellets were resuspended in 50 mL lysis buffer (20 mM Tris pH 8.0, 300 mM NaCl, 14.3 mM β-mercaptoethanol (BME), 1 mM EDTA pH 8.0, 5% (v/v) glycerol, 0.1% (v/v) Tween-20, 20 mM Imidazole pH 8.0) with a Pierce protease inhibitor tablet (Fisher PIA32955). The cells were lysed at 14,500 PSI by high-pressure homogenization with a microfluidizer (LM20 microfluidizer, Microfluidics). Phenylmethylsulfonyl fluoride (PMSF) at a final concentration of 1 uM was added immediately after lysis. Tagged hPUM1-HD and hPUM2-HD were isolated from the soluble fraction using NiNTA resin (Fisher PI88222). The collected resin was subsequently washed with 100 ml lysis buffer at a flow rate of 1 ml/min. Resin with tagged protein was reconstituted in 10 ml lysis buffer, then cleaved using 0.04% (v/v) 3C protease (Neta GSCRPT-Z03092-500) or ULP1 protease overnight. Recombinant protein was eluted by flow-through and then dialyzed into storage buffer (20 mM Tris pH 8.0, 100 mM NaCl, 1 mM DTT, 1 mM EDTA pH 8.0, 5% (v/v) glycerol, 0.1% (v/v) Tween-20). hPUM1-HD and hPUM2-HD were further purified on a Unosphere S column (BioRad) followed by a SEC70 column (BioRad) pre-equilibrated with storage buffer. Purified protein was then concentrated to 2-3 mg/ml.

### Molecular modeling system setup

In silico simulations were performed on published RNA-protein complex structures selected to match the in vitro screens performed: a) hPUM1-HD bound to UGUACAUC RNA for the canonical hPUM1-HD and RNA modification library screening (PDB ID: 3Q0P, (15)); b) hPUM1-HD bound to UGUAUAUA RNA for the canonical hPUM1-HD screening at site 8 (PDB ID: 3Q0N, (15)); c) hPUM2-HD bound to UGUAUAUA RNA for m^6^A screening (PDB ID: 3Q0Q, (15)). For the ψ screening, the 3Q0Q RNA structure was mutated from U to A at site 5 using Chimera (56) and energy minimized to acquire a structural model of hPUM2- HD bound to UGUAAAUA. A similar strategy was applied to 3Q0P to acquire a structural model of hPUM1-HD bound to UGUACAAU for the extended λD modified RNA screening. All hPUM1- HD and hPUM2-HD structures were optimized by PDB-REDO prior to use (57). Each system was set up similarly as in previous works (36). Briefly, the protein-nucleotide complexes were solvated in cubic boxes of explicit TIP3P water (58) using the CHARMM-GUI solution builder (59). Solvent padding of 10 Å was performed and a final ionic strength of 150 mM NaCl or KCl and 0-5 mM MgCl_2_ was used to match experimental conditions. Protonation states were applied based on PROPKA predictions at a pH of 7.0 (60). As mentioned in the results, both CHARMM36 and Amber force fields were used to represent protein and nucleic acid components (29,36,39,42,47,48). All systems were energy minimized prior to molecule dynamics to remove potential steric clashes.

### λ-dynamics calculations

A library of 48 bases, comprising 44 modified and 4 unmodified RNA candidates, were selected for screening with λD (36). Simulations were conducted using the CHARMM molecular simulation package (version c48b2) with the Basic λ-Dynamics Engine (BLaDE) for GPU accelerated modeling (37,38,61). Simulations were run at 25 °C and 1 atm, in the isothermal-isobaric ensemble. Long-range electrostatic interactions were modeled with particle mesh Ewald and Lennard-Jones interactions were truncated with force switching, with a cutoff of 10 Å and force switching beginning at 9 Å (62–64). SHAKE was used to constrain all bonds to hydrogen, facilitating the use of a 2 fs MD timestep (65). For purine-to-purine or pyrimidine-to-pyrimidine mutations, analogous atoms in the shared core were harmonically restrained to one another using the scaling of constrained atoms *(scat)* utility, developed for restraining alchemical substituents and protein backbones with λD (66,67).

Prior to production λD sampling, the Adaptive Landscape Flattening (ALF) algorithm (68,69) was used to identify optimal biasing potentials, which help facilitate frequent and even alchemical transitions between all perturbed molecules. For the initial modified RNA base screen with CHARMM force fields, 4 mutations per simulation per RNA site were performed, and mutations were grouped based on structural similarity. Optimal biases were identified after sampling ALF for a cumulative 48-55 ns, after which five independent production simulations were conducted for each group of modified RNA bases. Canonical base production simulations of hPUM1-HD and hPUM2-HD were run for 25-100 ns each. Convergence was confirmed by ensuring λD predictions remained unchanged within statistical noise as simulation times increased. For computing free energy differences, an initial 20% of the production data was excluded as equilibration. Additional extended modified RNA screenings were conducted in a pairwise manner with both CHARMM36 (39) and Amber force fields. In these head-to-head comparison simulations, production simulations were run for 50 ns each. Final ΔΔ*G*_bind_ values were then calculated by Boltzmann reweighting end state populations with WHAM (70) and computing statistical uncertainties with bootstrapping. The Amber ff14SB (41), OL3 (42), and modRNA08 (71) force fields were ported into the CHARMM simulation package using an open-source script developed in-house (https://github.com/murfalo/chamberr). This work was largely made possible by the open-source community’s work on the ParmEd parameter and topology editor (https://github.com/ParmEd/ParmEd). Previous work, which manually converted Amber ff14SB topologies and parameters into CHARMM-formatted files (41), was used as an initial test suite to validate the script-converted force fields.

### RNA oligonucleotide preparation

RNA oligonucleotides used for binding affinity experiments were synthesized on an ABI 394 DNA/RNA synthesizer (Applied Biosystems (ABI); Waltham, MA) as performed previously (36). m^6^A (10-3005-90; Glen Research; Sterling, VA), i^6^A (ANP-8615, Chemgene; Wilmington, MA), dhm^5^C (10-1510-95, Glen Research; Sterling, VA), m^5^C (10-3064- 90, Glen Research; Sterling, VA), and s^4^U (10-3052-90, Glen Research; Sterling, VA) (ANP-8626, Chemgene, Wilmington, MA) modified RNA phosphoramidites; fluorescein phosphoramidite (F5160, Lumiprobe; Cockeysville, MD); canonical DNA C phosphoramidite (ANP-5560, Chemgene; Wilmington, MA); and canonical RNA (A, ANP-5671; U, ANP-5674; C, ANP-6676; Chemgene, Wilmington, MA) phosphoramidites were purchased from commercial sources. Lacking commercial phosphoramidite reagents, ms^2^m^6^A-containing oligonucleotides were synthesized following previously established protocols (72–74). All synthesized oligos were gel extracted, lyophilized, and re-dissolved in water for downstream experiments. Concentrations of the aqueous RNA samples were determined by 260 nm UV absorption, using a Thermo Scientific Nanodrop One Spectrophotometer and theoretical 260 nm molar extinction coefficients provided by Integrated DNA Technologies.

### Electrophoretic mobility shift assay (EMSA)

Binding affinities of the synthesized RNA oligos to hPUM1-HD and hPUM2-HD were determined using EMSAs. Fluorescein-labeled RNA oligos were diluted in 1x EMSA binding buffer (10 mM HEPES pH 7.5, 50 mM KCl, 1 mM EDTA, 0.1% (v/v) Tween-20, 1 mM DTT, 0.01 mg/ml BSA (BP9706100; Fisher Scientific; Hampton NH)), to a 10x concentration of 20 and 50 nM for hPUM1 and hPUM2 experiments, respectively. Serial protein dilutions were prepared for a final volume of 10 uL as follows. In 1x EMSA binding buffer, proteins were diluted to an initial 1.1x concentration of 1111 nM and three-fold serial diluted. In tubes, 9 ul protein and 1 ul RNA were mixed together for final protein concentrations of 1000 nM, 333 nM, 111 nM, 37 nM, 12.3 nM, 4.1 nM, 1.4 nM with 2 and 5 nM RNA for hPUM1 and hPUM2 experiments, respectively. An eighth lane with no protein was prepared using EMSA binding buffer alone. After incubating the protein-RNA sample at 4°C for 30 minutes, 2 uL of 6x EMSA loading buffer (15% (w/v) Ficoll 400, 0.01% (w/v) Bromophenol Blue) was added to each 10 uL dilution for gel loading. Non-denaturing, polyacrylamide gels (0.5x TBE (Tris/Borate/EDTA buffer; 45 mM Tris, 45 mM boric acid, 1 mM EDTA), 5% (v/v) Acrylamide/Bis 19:1; catalyzed with 0.1% (w/v) ammonium persulfate (APS) and 0.1% (v/v) N,N,N′,N′-Tetramethylethylenediamine (TEMED)) were pre-equilibrated in 0.5x TBE using a Mini-PROTEAN® vertical electrophoresis cell (Bio Rad; 1658005) for 30 minutes at 4°C. Samples were loaded and run at 75 volts for 45 minutes at 4°C. The gels were imaged with a Bio Rad ChemiDoc imager on a U/V tray capable of measuring Flourescein 528/532 nm absorption. Flourescein absorption intensities of the lower (unbound) and upper (bound) RNA bands were quantified using Fiji ImageJ (75). K_d_ values were calculated with a non-linear fit in GraphPad Prism version 10.1.1 for MacOS (GraphPad Software, Boston, MA). Averages, standard deviations, and graphs were also performed and made in GraphPad Prism. All EMSA experiments were performed in a minimum of three replicates, with additional replicates conducted as necessary to ensure that the standard error of the mean (SEM) for all estimated K_d_ values remained below 10. Statistical significance of differences in K_d_ between oligos were obtained using one-way ANOVA.

### Data collection & analysis

To complement experimental EMSA binding data, additional hPUM1 and hPUM2 RNA in vitro binding data was collected from the literature (18,26). For unmodified RNAs, matching sequences and their variations to this study’s binding data were selected from previously published data (18). Duplicate entries were averaged into a single experimental Δ*G* for each sequence. In total, experimental Δ*G*s were obtained for 21 of 24 possible single-site mutants of UGUAUAUA bound to hPUM2. For hPUM1, experimental Δ*G*s were obtained for 8 mutants of UGUACAUC. Binding data for 6 Ψ and m^6^A mutants to hPUM2 were obtained from (26).

For comparison of λD predictions with experimental values, experimental K_d_s were first converted to absolute Δ*G*s using the relationship Δ*G* = −*RT* ln *K_d_* at the experimental temperature. Next, λD predicted Δ*G*s were converted to absolute Δ*G*s following established methods (76). Briefly, the data was first divided into groups based on the consensus oligo and protein target. For each dataset, the mean experimental Δ*G* (Δ*G̅_expt_*) and predicted in silico Δ*G* (Δ*G̅_pred_*) were calculated. Finally, the in silico Δ*G_pred_* were converted to absolute Δ*G_pred_* by recentering values relative to the experimental mean Δ*G̅_expt_*. In sum, Δ*G_pred_* = Δ*G_pred_* − (Δ*G̅_pred_* − Δ*G̅_expt_*) for each Δ*G_pred_* in the dataset (76). The λD predicted Δ*G*s (Δ*G_pred_*) and experimental Δ*G*s (Δ*G_pred_*) were then directly compared.

## Acknowledgements

The authors thank members of the Aoki, Vilseck, and Zhang Labs for their helpful discussion and acknowledge the Indiana University Pervasive Technology Institute for providing supercomputing and storage resources that have contributed to the research results reported within this paper. S.T.A, W.Z., and J.Z.V. received start-up funds from the Indiana University School of Medicine and its Precision Health Initiative (PHI). J.Z.V. is funded by the National Institute of General Medical Sciences (NIGMS) of the National Institutes of Health (NIH) (R35GM146888). S.T.A. is funded by the NIH/NIGMS (R35GM142691) and an Indiana University Research Support Funds Grant (RSFG). E.K. is funded by UMO-2020/39/B/NZ1/03054 and R.K funded by UMO- 2022/45/B/ST4/03586.

UMO-2020/39/B/NZ1/03054 to EK and UMO-2022/45/B/ST4/03586 to RK

## Supplemental Figure captions

**Table 1**. Human Pumilio 1 and 2 λ-dynamics results to unmodified RNA.

**Table 2**. Human Pumilio 1 λ-dynamics results to modified RNA.

**Table 3**. Human Pumilio 2 λ-dynamics and in vitro binding results to modified RNA.

**Fig S1.** ANOVA binding statistics. Binding statistics of **(A)** human Pumilio 1 (hPUM1) and **(B)** human Pumilio 2 (hPUM2) in vitro binding results to wild type or modified RNA. All results were compared to wild type RNA. See methods for further details.

**Fig S2.** Human Pumilio 2 binding to modified RNAs in vitro was also consistent to λ-dynamics predictions in silico. **(A)** Electrophoretic Mobility Shift Assays (EMSAs) were used to estimate the binding affinity in vitro. Synthetic RNA oligos without or with the designated RNA modifications were incubated with increasing concentrations (0-1000 nM) of recombinant human Pumilio 2 (hPUM2) RNA binding domain, run on a polyacrylamide gel, and imaged for carboxyfluorescein (FAM) fluorescence. A sample without protein served as an unbound RNA control. The lower band corresponds to unbound RNA. The upper band corresponds to RNA bound to the recombinant protein. The binding affinity can be estimated by calculating the protein concentration at the binding inflection point. Shown are representative gels from hPUM2 EMSA binding experiments. **(B)** EMSAs were performed at least three times, and calculated binding dissociation constants (K_d_) reported with their mean and standard deviation. *, p <0.05; **, p <0.01; ***, p <0.001. See **Fig S1** for full statistical analyses and the methods for more experimental details.

---

**Figure S1.**
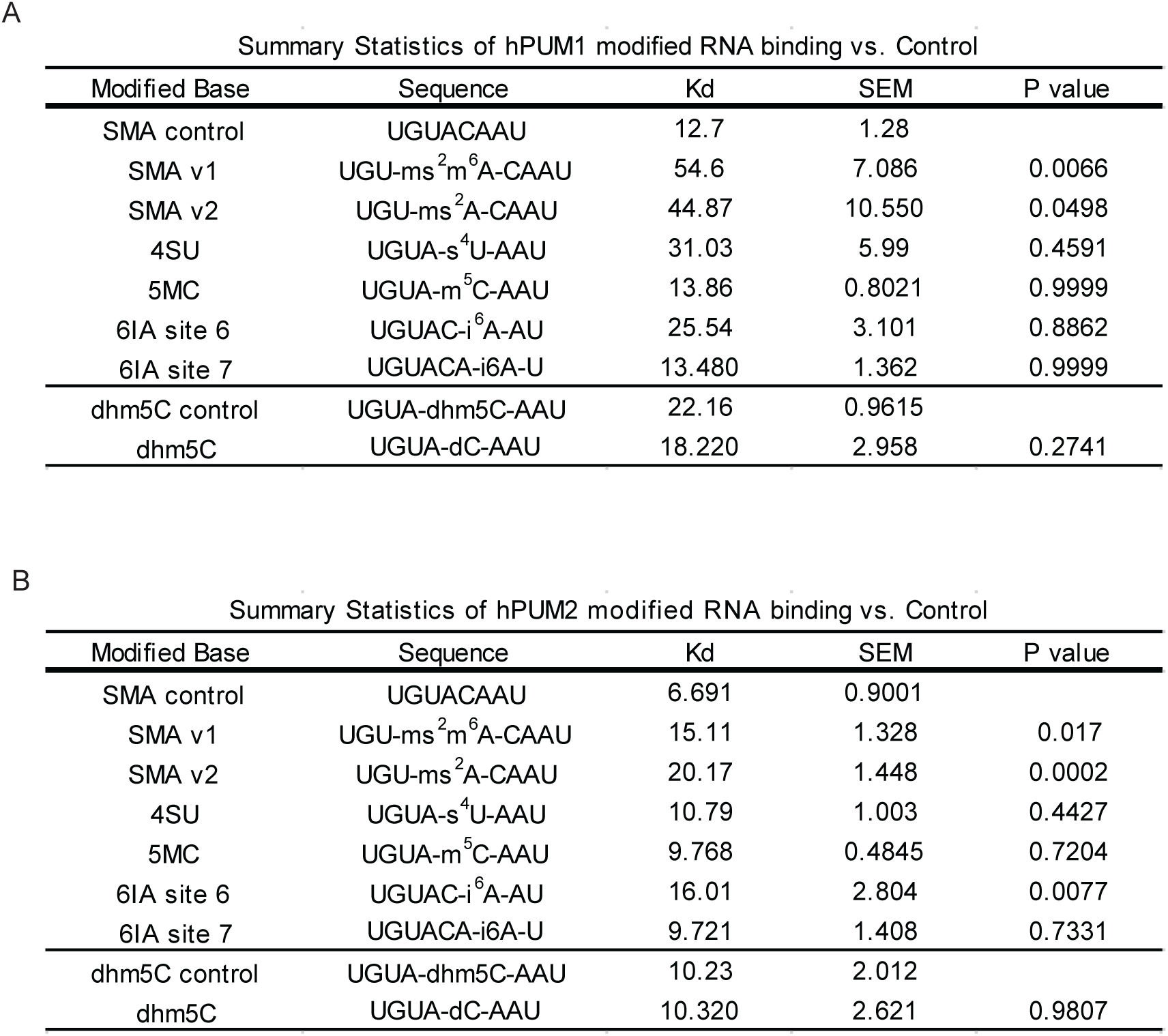

**Table S1.**
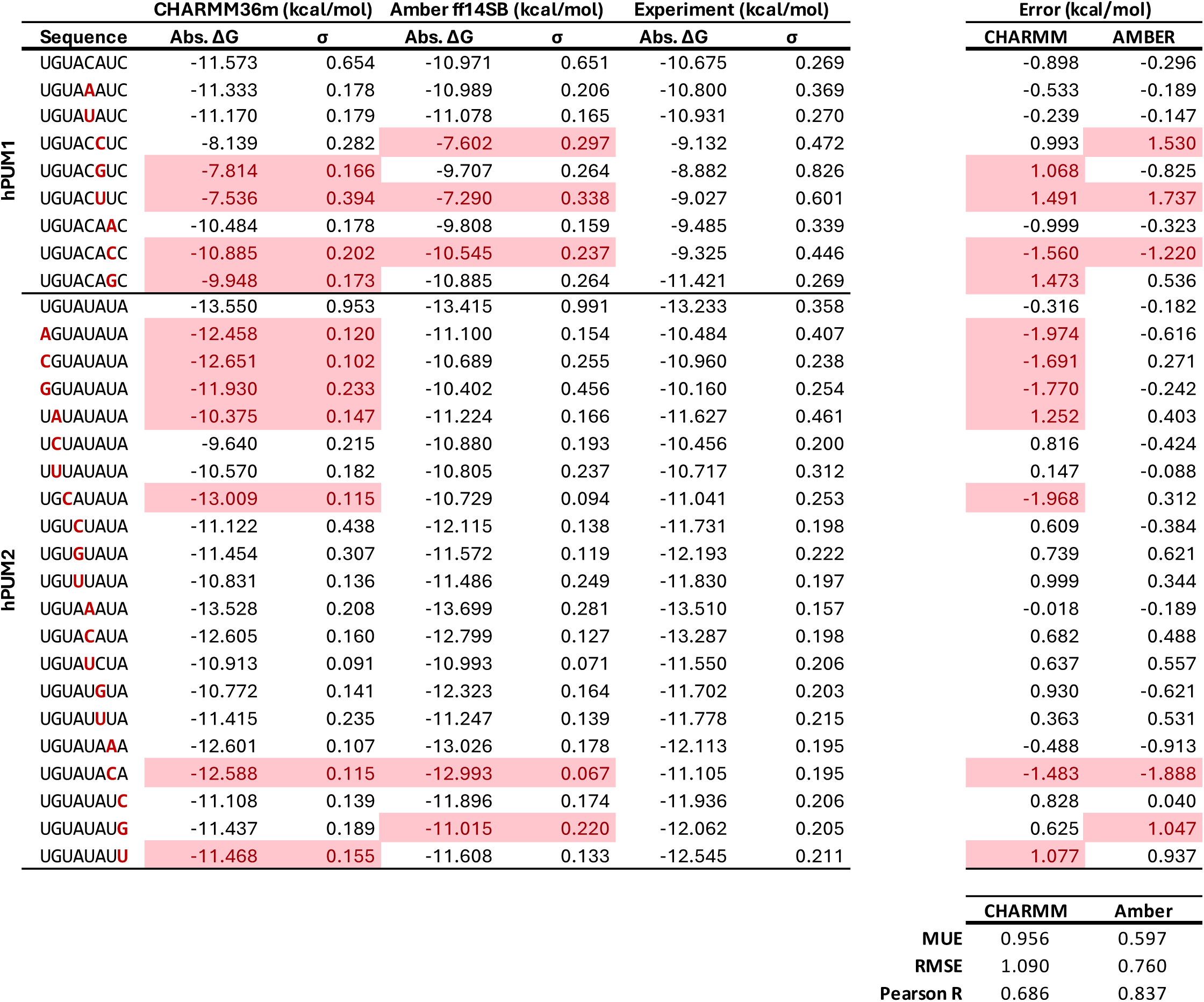

**Table S2.**
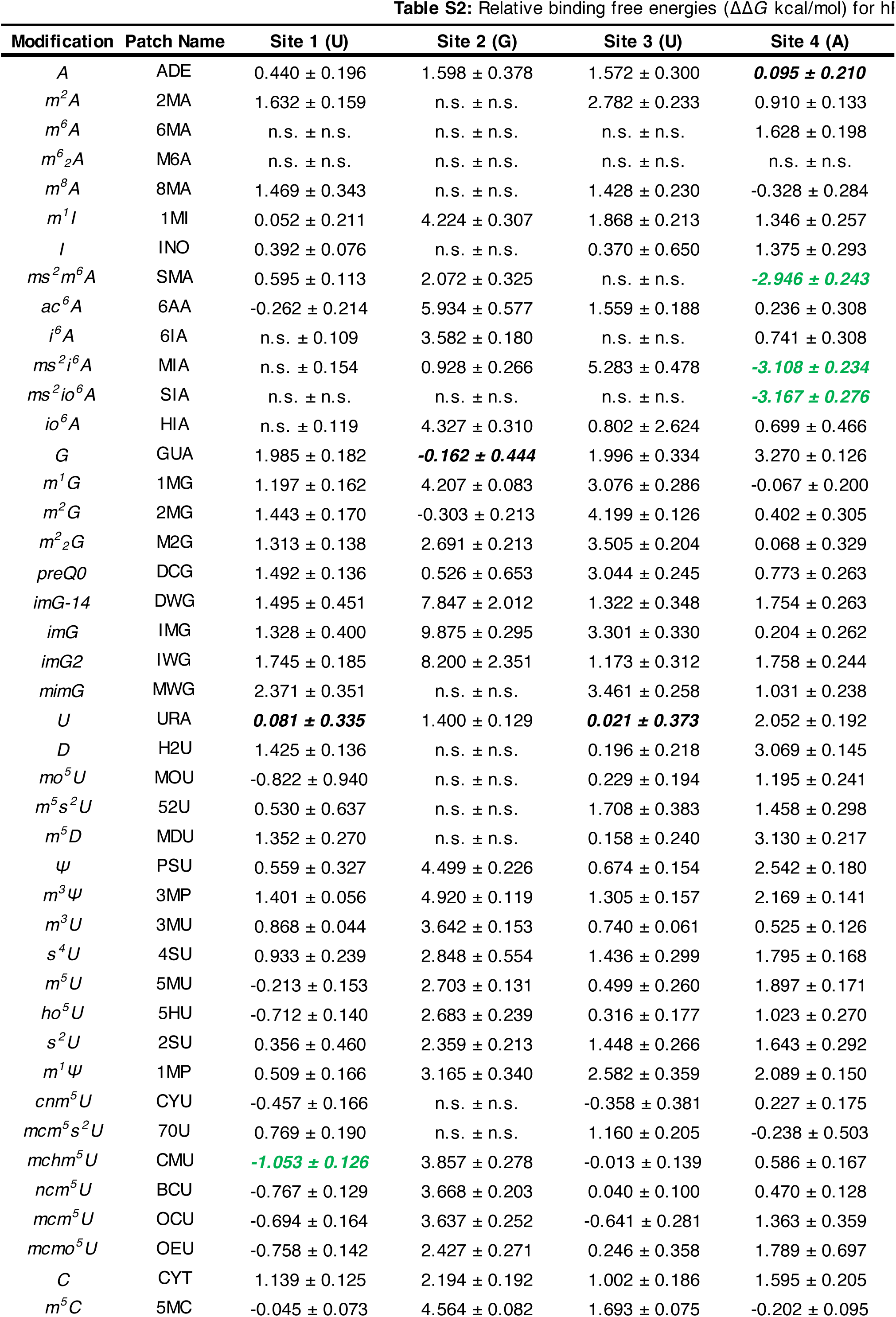
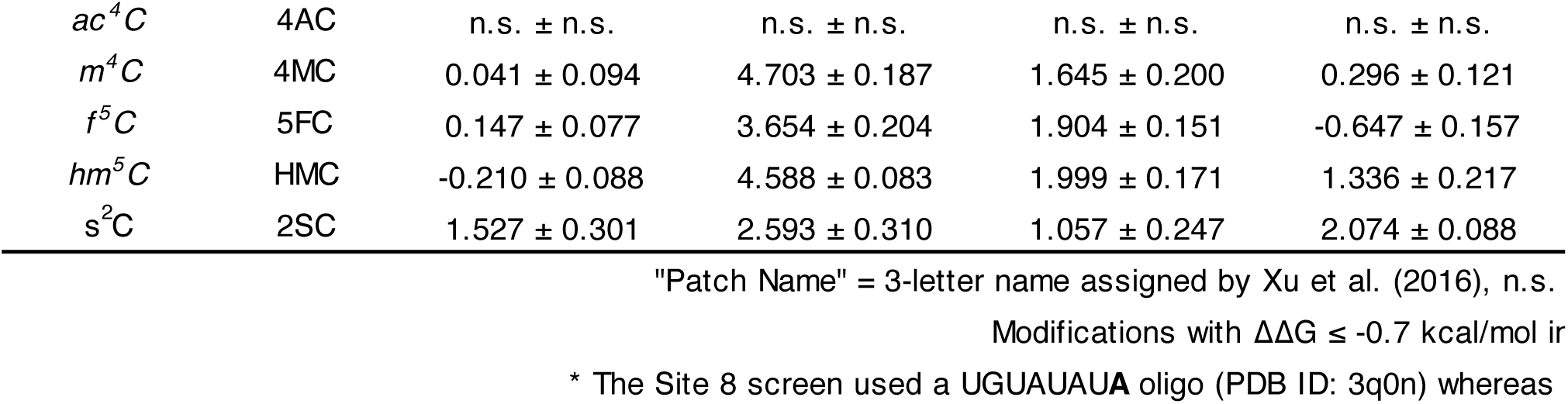
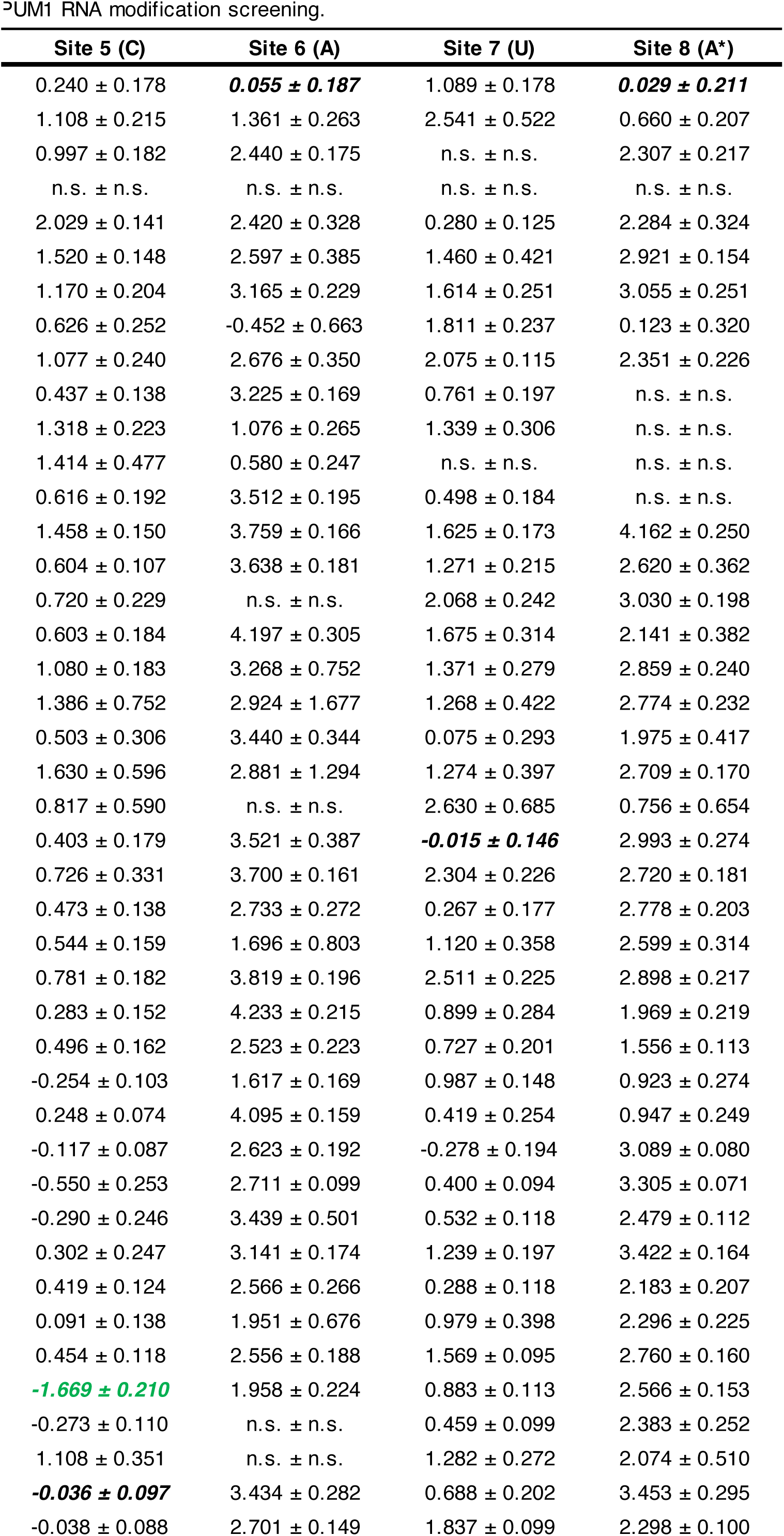
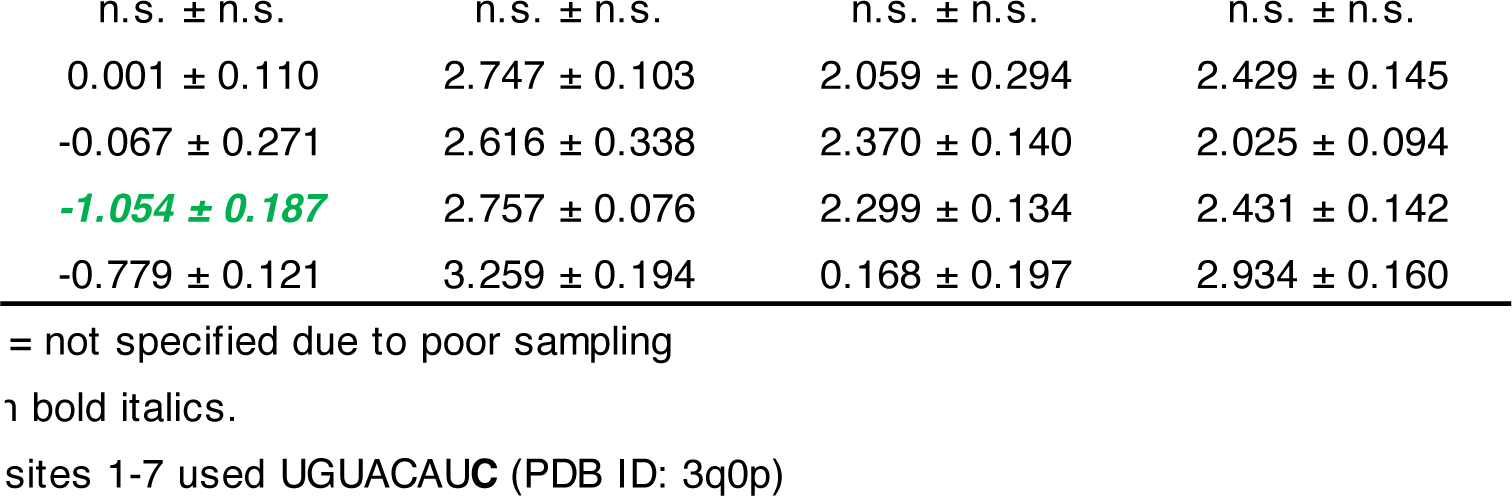

| Modification | Patch Name | Site 1 (U) | Site 2 (G) | Site 3 (U) | Site 4 (A) |
| --- | --- | --- | --- | --- | --- |
| A | ADE | 0.440 ± 0.196 | 1.598 ± 0.378 | 1.572 ± 0.300 | **0.095 ± 0.210** |
| $m^2A$ | 2MA | 1.632 ± 0.159 | n.s. ± n.s. | 2.782 ± 0.233 | 0.910 ± 0.133 |
| $m^6A$ | 6MA | n.s. ± n.s. | n.s. ± n.s. | n.s. ± n.s. | 1.628 ± 0.198 |
| $m^6_2A$ | M6A | n.s. ± n.s. | n.s. ± n.s. | n.s. ± n.s. | n.s. ± n.s. |
| $m^8A$ | 8MA | 1.469 ± 0.343 | n.s. ± n.s. | 1.428 ± 0.230 | -0.328 ± 0.284 |
| $m^1I$ | 1MI | 0.052 ± 0.211 | 4.224 ± 0.307 | 1.868 ± 0.213 | 1.346 ± 0.257 |
| I | INO | 0.392 ± 0.076 | n.s. ± n.s. | 0.370 ± 0.650 | 1.375 ± 0.293 |
| $ms^2m^6A$ | SMA | 0.595 ± 0.113 | 2.072 ± 0.325 | n.s. ± n.s. | **-2.946 ± 0.243** |
| $ac^6A$ | 6AA | -0.262 ± 0.214 | 5.934 ± 0.577 | 1.559 ± 0.188 | 0.236 ± 0.308 |
| $i^6A$ | 6IA | n.s. ± 0.109 | 3.582 ± 0.180 | n.s. ± n.s. | 0.741 ± 0.308 |
| $ms^2i^6A$ | MIA | n.s. ± 0.154 | 0.928 ± 0.266 | 5.283 ± 0.478 | **-3.108 ± 0.234** |
| $ms^2io^6A$ | SIA | n.s. ± n.s. | n.s. ± n.s. | n.s. ± n.s. | **-3.167 ± 0.276** |
| $io^6A$ | HIA | n.s. ± 0.119 | 4.327 ± 0.310 | 0.802 ± 2.624 | 0.699 ± 0.466 |
| G | GUA | 1.985 ± 0.182 | **-0.162 ± 0.444** | 1.996 ± 0.334 | 3.270 ± 0.126 |
| $m^1G$ | 1MG | 1.197 ± 0.162 | 4.207 ± 0.083 | 3.076 ± 0.286 | -0.067 ± 0.200 |
| $m^2G$ | 2MG | 1.443 ± 0.170 | -0.303 ± 0.213 | 4.199 ± 0.126 | 0.402 ± 0.305 |
| $m^2_2G$ | M2G | 1.313 ± 0.138 | 2.691 ± 0.213 | 3.505 ± 0.204 | 0.068 ± 0.329 |
| preQ0 | DCG | 1.492 ± 0.136 | 0.526 ± 0.653 | 3.044 ± 0.245 | 0.773 ± 0.263 |
| imG-14 | DWG | 1.495 ± 0.451 | 7.847 ± 2.012 | 1.322 ± 0.348 | 1.754 ± 0.263 |
| imG | IMG | 1.328 ± 0.400 | 9.875 ± 0.295 | 3.301 ± 0.330 | 0.204 ± 0.262 |
| imG2 | IWG | 1.745 ± 0.185 | 8.200 ± 2.351 | 1.173 ± 0.312 | 1.758 ± 0.244 |
| mimG | MWG | 2.371 ± 0.351 | n.s. ± n.s. | 3.461 ± 0.258 | 1.031 ± 0.238 |
| U | URA | **0.081 ± 0.335** | 1.400 ± 0.129 | **0.021 ± 0.373** | 2.052 ± 0.192 |
| D | H2U | 1.425 ± 0.136 | n.s. ± n.s. | 0.196 ± 0.218 | 3.069 ± 0.145 |
| $mo^5U$ | MOU | -0.822 ± 0.940 | n.s. ± n.s. | 0.229 ± 0.194 | 1.195 ± 0.241 |
| $m^5s^2U$ | 52U | 0.530 ± 0.637 | n.s. ± n.s. | 1.708 ± 0.383 | 1.458 ± 0.298 |
| $m^5D$ | MDU | 1.352 ± 0.270 | n.s. ± n.s. | 0.158 ± 0.240 | 3.130 ± 0.217 |
| $\psi$ | PSU | 0.559 ± 0.327 | 4.499 ± 0.226 | 0.674 ± 0.154 | 2.542 ± 0.180 |
| $m^3\psi$ | 3MP | 1.401 ± 0.056 | 4.920 ± 0.119 | 1.305 ± 0.157 | 2.169 ± 0.141 |
| $m^3U$ | 3MU | 0.868 ± 0.044 | 3.642 ± 0.153 | 0.740 ± 0.061 | 0.525 ± 0.126 |
| $s^4U$ | 4SU | 0.933 ± 0.239 | 2.848 ± 0.554 | 1.436 ± 0.299 | 1.795 ± 0.168 |
| $m^5U$ | 5MU | -0.213 ± 0.153 | 2.703 ± 0.131 | 0.499 ± 0.260 | 1.897 ± 0.171 |
| $ho^5U$ | 5HU | -0.712 ± 0.140 | 2.683 ± 0.239 | 0.316 ± 0.177 | 1.023 ± 0.270 |
| $s^2U$ | 2SU | 0.356 ± 0.460 | 2.359 ± 0.213 | 1.448 ± 0.266 | 1.643 ± 0.292 |
| $m^1\psi$ | 1MP | 0.509 ± 0.166 | 3.165 ± 0.340 | 2.582 ± 0.359 | 2.089 ± 0.150 |
| $cnm^5U$ | CYU | -0.457 ± 0.166 | n.s. ± n.s. | -0.358 ± 0.381 | 0.227 ± 0.175 |
| $mcm^5s^2U$ | 70U | 0.769 ± 0.190 | n.s. ± n.s. | 1.160 ± 0.205 | -0.238 ± 0.503 |
| $mchm^5U$ | CMU | **-1.053 ± 0.126** | 3.857 ± 0.278 | -0.013 ± 0.139 | 0.586 ± 0.167 |
| $ncm^5U$ | BCU | -0.767 ± 0.129 | 3.668 ± 0.203 | 0.040 ± 0.100 | 0.470 ± 0.128 |
| $mcm^5U$ | OCU | -0.694 ± 0.164 | 3.637 ± 0.252 | -0.641 ± 0.281 | 1.363 ± 0.359 |
| $mcmo^5U$ | OEU | -0.758 ± 0.142 | 2.427 ± 0.271 | 0.246 ± 0.358 | 1.789 ± 0.697 |
| C | CYT | 1.139 ± 0.125 | 2.194 ± 0.192 | 1.002 ± 0.186 | 1.595 ± 0.205 |
| $m^5C$ | 5MC | -0.045 ± 0.073 | 4.564 ± 0.082 | 1.693 ± 0.075 | -0.202 ± 0.095 |
| $ac^4C$ | 4AC | n.s. ± n.s. | n.s. ± n.s. | n.s. ± n.s. | n.s. ± n.s. |
| $m^4C$ | 4MC | $0.041 \pm 0.094$ | $4.703 \pm 0.187$ | $1.645 \pm 0.200$ | $0.296 \pm 0.121$ |
| $f^5C$ | 5FC | $0.147 \pm 0.077$ | $3.654 \pm 0.204$ | $1.904 \pm 0.151$ | $-0.647 \pm 0.157$ |
| $hm^5C$ | HMC | $-0.210 \pm 0.088$ | $4.588 \pm 0.083$ | $1.999 \pm 0.171$ | $1.336 \pm 0.217$ |
| $s^2C$ | 2SC | $1.527 \pm 0.301$ | $2.593 \pm 0.310$ | $1.057 \pm 0.247$ | $2.074 \pm 0.088$ |

| Site 5 (C) | Site 6 (A) | Site 7 (U) | Site 8 (A*) |
| --- | --- | --- | --- |
| 0.240 ± 0.178 | **0.055 ± 0.187** | 1.089 ± 0.178 | **0.029 ± 0.211** |
| 1.108 ± 0.215 | 1.361 ± 0.263 | 2.541 ± 0.522 | 0.660 ± 0.207 |
| 0.997 ± 0.182 | 2.440 ± 0.175 | n.s. ± n.s. | 2.307 ± 0.217 |
| n.s. ± n.s. | n.s. ± n.s. | n.s. ± n.s. | n.s. ± n.s. |
| 2.029 ± 0.141 | 2.420 ± 0.328 | 0.280 ± 0.125 | 2.284 ± 0.324 |
| 1.520 ± 0.148 | 2.597 ± 0.385 | 1.460 ± 0.421 | 2.921 ± 0.154 |
| 1.170 ± 0.204 | 3.165 ± 0.229 | 1.614 ± 0.251 | 3.055 ± 0.251 |
| 0.626 ± 0.252 | -0.452 ± 0.663 | 1.811 ± 0.237 | 0.123 ± 0.320 |
| 1.077 ± 0.240 | 2.676 ± 0.350 | 2.075 ± 0.115 | 2.351 ± 0.226 |
| 0.437 ± 0.138 | 3.225 ± 0.169 | 0.761 ± 0.197 | n.s. ± n.s. |
| 1.318 ± 0.223 | 1.076 ± 0.265 | 1.339 ± 0.306 | n.s. ± n.s. |
| 1.414 ± 0.477 | 0.580 ± 0.247 | n.s. ± n.s. | n.s. ± n.s. |
| 0.616 ± 0.192 | 3.512 ± 0.195 | 0.498 ± 0.184 | n.s. ± n.s. |
| 1.458 ± 0.150 | 3.759 ± 0.166 | 1.625 ± 0.173 | 4.162 ± 0.250 |
| 0.604 ± 0.107 | 3.638 ± 0.181 | 1.271 ± 0.215 | 2.620 ± 0.362 |
| 0.720 ± 0.229 | n.s. ± n.s. | 2.068 ± 0.242 | 3.030 ± 0.198 |
| 0.603 ± 0.184 | 4.197 ± 0.305 | 1.675 ± 0.314 | 2.141 ± 0.382 |
| 1.080 ± 0.183 | 3.268 ± 0.752 | 1.371 ± 0.279 | 2.859 ± 0.240 |
| 1.386 ± 0.752 | 2.924 ± 1.677 | 1.268 ± 0.422 | 2.774 ± 0.232 |
| 0.503 ± 0.306 | 3.440 ± 0.344 | 0.075 ± 0.293 | 1.975 ± 0.417 |
| 1.630 ± 0.596 | 2.881 ± 1.294 | 1.274 ± 0.397 | 2.709 ± 0.170 |
| 0.817 ± 0.590 | n.s. ± n.s. | 2.630 ± 0.685 | 0.756 ± 0.654 |
| 0.403 ± 0.179 | 3.521 ± 0.387 | **-0.015 ± 0.146** | 2.993 ± 0.274 |
| 0.726 ± 0.331 | 3.700 ± 0.161 | 2.304 ± 0.226 | 2.720 ± 0.181 |
| 0.473 ± 0.138 | 2.733 ± 0.272 | 0.267 ± 0.177 | 2.778 ± 0.203 |
| 0.544 ± 0.159 | 1.696 ± 0.803 | 1.120 ± 0.358 | 2.599 ± 0.314 |
| 0.781 ± 0.182 | 3.819 ± 0.196 | 2.511 ± 0.225 | 2.898 ± 0.217 |
| 0.283 ± 0.152 | 4.233 ± 0.215 | 0.899 ± 0.284 | 1.969 ± 0.219 |
| 0.496 ± 0.162 | 2.523 ± 0.223 | 0.727 ± 0.201 | 1.556 ± 0.113 |
| -0.254 ± 0.103 | 1.617 ± 0.169 | 0.987 ± 0.148 | 0.923 ± 0.274 |
| 0.248 ± 0.074 | 4.095 ± 0.159 | 0.419 ± 0.254 | 0.947 ± 0.249 |
| -0.117 ± 0.087 | 2.623 ± 0.192 | -0.278 ± 0.194 | 3.089 ± 0.080 |
| -0.550 ± 0.253 | 2.711 ± 0.099 | 0.400 ± 0.094 | 3.305 ± 0.071 |
| -0.290 ± 0.246 | 3.439 ± 0.501 | 0.532 ± 0.118 | 2.479 ± 0.112 |
| 0.302 ± 0.247 | 3.141 ± 0.174 | 1.239 ± 0.197 | 3.422 ± 0.164 |
| 0.419 ± 0.124 | 2.566 ± 0.266 | 0.288 ± 0.118 | 2.183 ± 0.207 |
| 0.091 ± 0.138 | 1.951 ± 0.676 | 0.979 ± 0.398 | 2.296 ± 0.225 |
| 0.454 ± 0.118 | 2.556 ± 0.188 | 1.569 ± 0.095 | 2.760 ± 0.160 |
| **-1.669 ± 0.210** | 1.958 ± 0.224 | 0.883 ± 0.113 | 2.566 ± 0.153 |
| -0.273 ± 0.110 | n.s. ± n.s. | 0.459 ± 0.099 | 2.383 ± 0.252 |
| 1.108 ± 0.351 | n.s. ± n.s. | 1.282 ± 0.272 | 2.074 ± 0.510 |
| **-0.036 ± 0.097** | 3.434 ± 0.282 | 0.688 ± 0.202 | 3.453 ± 0.295 |
| -0.038 ± 0.088 | 2.701 ± 0.149 | 1.837 ± 0.099 | 2.298 ± 0.100 |

|  |  |  |  |
| --- | --- | --- | --- |
| n.s. ± n.s. | n.s. ± n.s. | n.s. ± n.s. | n.s. ± n.s. |
| 0.001 ± 0.110 | 2.747 ± 0.103 | 2.059 ± 0.294 | 2.429 ± 0.145 |
| -0.067 ± 0.271 | 2.616 ± 0.338 | 2.370 ± 0.140 | 2.025 ± 0.094 |
| ***-1.054 ± 0.187*** | 2.757 ± 0.076 | 2.299 ± 0.134 | 2.431 ± 0.142 |
| -0.779 ± 0.121 | 3.259 ± 0.194 | 0.168 ± 0.197 | 2.934 ± 0.160 |

= not specified due to poor sampling
1 bold italics.
sites 1-7 used UGUACAUC (PDB ID: 3q0p)

**Table S3.**

|  | Sequence | CHARMM36m (kcal/mol) |  | Amber modrna08 (kcal/mol) |  | Experiment (kcal/mol) |  | Error (kcal/mol) |  |
| --- | --- | --- | --- | --- | --- | --- | --- | --- | --- |
| | | $\Delta G$ | $\sigma$ | $\Delta G$ | $\sigma$ | $\Delta G$ | $\sigma$ | CHARMM | AMBER |
| $\psi$ | CCUGUAAAUA | -13.402 | 0.235 | -12.204 | 0.319 | -12.510 | 0.045 | -0.892 | -0.306 |
| | CCΨGUAAAUA | -12.976 | 0.094 | -12.005 | 0.033 | -12.181 | 0.026 | -0.794 | -0.176 |
| | CCUGΨAAAUA | -12.223 | 0.132 | -13.137 | 0.270 | -12.024 | 0.118 | -0.199 | 1.113 |
| | CCUGUAAAΨA | -11.923 | 0.170 | -11.978 | 0.166 | -11.916 | 0.066 | -0.007 | 0.062 |
| $m^6A$ | CCUGUAUAUA | -15.094 | 0.215 | -14.022 | 0.291 | -13.160 | 0.000 | -1.934 | 0.862 |
| | CCUGU$AUAUA$ | -13.337 | 0.166 | -13.303 | 0.151 | -13.061 | 0.046 | -0.276 | 0.242 |
| | CCUGUAU$AUA$ | -13.204 | 0.082 | -12.436 | 0.231 | -12.597 | 0.031 | -0.608 | -0.161 |
| | CCUGUAUAU$A$ | -12.651 | 0.110 | -13.455 | 0.093 | -12.547 | 0.258 | -0.105 | 0.908 |
| PUM modRNAs | UGUACAAU | -10.621 | 0.383 | -11.356 | 0.360 | -10.691 | 0.116 | 0.070 | 0.665 |
| | UGU-$SMA$-CAAU | -13.142 | 0.228 | -10.296 | 0.162 | -9.897 | 0.203 | -3.245 | 0.400 |
| | UGUA-$4SU$-AAU | -10.562 | 0.128 | -10.011 | 0.160 | -10.231 | 0.302 | -0.331 | -0.220 |
| | UGUA-$5MC$-AAU | -10.891 | 0.074 | -10.706 | 0.134 | -10.631 | 0.130 | -0.260 | 0.075 |
| | UGUA-$dHMC$-AAU | -11.125 | 0.197 | -10.608 | 0.094 | -10.248 | 0.134 | -0.877 | 0.360 |
| | UGUAC-$6IA$-AU | -6.767 | 0.099 | -9.951 | 0.168 | -10.653 | 0.112 | 3.886 | -0.701 |
| | UGUACA-$6IA$-U | -9.801 | 0.155 | -10.715 | 0.151 | -10.469 | 0.137 | 0.668 | 0.246 |
|  |  | CHARMM |  | Amber |  |  |  |  |  |
| MUE |  | 0.943 |  | 0.433 |  |  |  |  |  |
| RMSE |  | 1.476 |  | 0.539 |  |  |  |  |  |
| Pearson R |  | 0.669 |  | 0.929 |  |  |  |  |  |

